# Machine learning inference of natural product chemistry across biosynthetic gene cluster types

**DOI:** 10.1101/2025.03.13.642868

**Authors:** Martin Larralde, Georg Zeller

## Abstract

With ever-increasing volumes of sequencing data for biosynthetic gene clusters (BGCs), computational methods for the prediction of resulting secondary metabolites are critically needed. Here, we present CHAMOIS, a machine learning tool inferring metabolite properties from protein domains in BGCs. Out of 539 relevant chemical properties from the ChemOnt ontology, CHAMOIS predicts 120 with an AUPRC > 0.5. Although entirely data-driven, CHAMOIS infers many protein-metabolite links that are consistent with the scientific literature and suggests interesting novel biosynthetic functions of uncharacterized proteins. Finally, to guide experimental BGC characterisation, CHAMOIS can pinpoint which BGC within a given genome produces a pre-specified metabolite.

## Background

Microorganisms across Earth’s habitats are capable of producing an astonishing diversity of natural products. The enzymatic machinery for synthesising many of these compounds is encoded by genes located in genomic proximity and thus referred to as biosynthetic gene clusters (BGCs). BGC-encoded enzymes often interact in biosynthetic pathways in a modular fashion. Evolution acting on these genetic building blocks has given rise to a vast diversity of cluster architectures and a resulting biochemical diversity of produced molecules(1). While BGCs usually contain a core biosynthetic cluster composed of key enzymes(2), these are often surrounded by additional genes involved in regulation or metabolite transport(3).

Historically, BGCs were often studied in microbial isolates known to synthesize a secondary metabolite of interest. BGC identification usually relied on genetic techniques such as cosmid library generation(4) or knock-out studies targeting putative biosynthetic or resistance genes(5). More recently, the deluge of genomic data warranted the development of novel methods for identifying candidate BGCs *in silico*. Several such methods are based on rules derived from expert knowledge, including the popular antiSMASH software(6), which combines the detection of key biosynthetic genes with a system of biochemistry-aware rules. In parallel, standardised repositories of experimentally-validated BGCs, such as MIBiG(7), encouraged the development of a new generation of machine learning (ML) and artificial intelligence (AI) methods to complement rule-based methods. These include ClusterFinder(8), DeepBGC(9), GECCO(10) and SanntiS(11).

With an ever-increasing amount of genomic data, the sprawl of BGC prediction methods has led to an exponential accumulation of BGC predictions: the first large-scale repository of BGC prediction, the antiSMASH-db (2017), contained ∼22,000 BGC predictions made by antiSMASH on ∼4,000 bacterial genomes(12); six years later, the proGenomes3 database (2023) released more than 3 million BGC predictions made with GECCO(10,13). The most comprehensive resource for experimentally-validated BGCs, MIBiG 4.0 (2024), however, numbers only 2,437 “active” entries (https://mibig.secondarymetabolites.org/stats) . Hence, for the vast majority of *in silico* predicted microbial BGCs, the encoded metabolite is unknown, since natural products remain more challenging to identify than the genomic locations of BGCs. Only BGCs with strong homology to experimentally characterized ones can be confidently hypothesised to produce a similar metabolite. However, even the deletion of small accessory genes can substantially alter the final product, its biochemical properties and biological activity(14). BGCs with remote homology to known instances are very challenging to characterise *in silico*(15). Even today, natural products discovery from genomic data is limited by time-consuming experiments, and the large volumes of available BGC predictions remain relatively unexplored, underlining the need for better computational metabolite inference and prioritisation methods.

Toward this goal, antiSMASH categorises BGCs according to one or more high-level product types based on its ruleset (101 distinct types in v8.0). These types summarise the broad biosynthetic pathway, but often reveal little detail about the final molecule itself. Currently available data-driven approaches for BGC categorisation assign BGCs to one or several of six more coarse-grained MIBiG types, typically using relatively simple ML classifiers, as e.g. implemented in DeepBGC(9). While the predicted types can be useful for prioritising BGCs for experimentation, their information content is generally too low to derive interesting chemical properties or bioactivities of the putative product.

Among the different types of BGCs, relationship between enzyme sequence and corresponding metabolite properties have been mostly studied for the polyketide (PK) and non-ribosomal peptide (NRP) types, as they often feature multifunctional enzymes composed of multiple biosynthetic domains. These domains form an assembly line and incorporate smaller precursors into the molecule, which is modified sequentially to form a larger metabolite backbone(16). Several studies have shown the specificity of certain PK or NRP synthase domains for particular precursors based on sequence features(17,18). These discoveries have in turn been integrated into computational methods for inferring some aspects of natural product biosynthesis, such as NRPSpredictor2(19), NERPA(20), RiPPMiner(21), transAtor(22), or PRISM(23), the three latter of which can be used to predict the backbone structure of BGC products from sequence features, but only for certain BGC types. While these methods represent a step towards natural product prediction for certain BGCs, consistent and accurate product prediction across all BGC types remains a fundamental challenge, despite the elucidation of more and more biosynthetic pathways.

Here, to approach this problem from a novel data-driven perspective, we developed CHAMOIS, the first method to employ machine-learning for predicting chemical properties of BGCs products from gene sequences across all major types of BGCs. We evaluate its prediction accuracy in carefully stratified cross-validation to assess how CHAMOIS generalizes across distinct BGCs and metabolites. We further demonstrate that CHAMOIS’s data-driven model captures relevant known and novel biochemical information about enzymatic domains, and finally show that CHAMOIS is useful to pinpoint which BGC within a given genome gives rise to an a priori known metabolite – a capability that could expedite experimental characterisation of BGCs and discovery of novel biosynthetic enzymes.

## Results

### CHAMOIS: A novel machine-learning method for prediction of chemical properties from a BGC sequence

Inferring chemical structures directly is nearly impossible because of the sheer size of chemical space. The limited availability of experimentally validated BGCs for training and testing poses an additional challenge which renders the task of directly inferring metabolite structures from BGC sequences extremely difficult. Therefore, instead of generating candidate metabolite structures, we developed a method for predicting chemical properties of these metabolites using machine learning to infer links between gene sequences and chemical properties from experimentally characterized BGCs with known metabolites.

To capture chemical properties of arbitrary natural products for ML inference, we use the classes of a chemical ontology, ChemOnt, which contains 4,825 non-exclusive chemical categories covering all domains of chemistry(24). This representation of molecules into a lower-dimensional space of relevant properties resembles molecular fingerprinting, a common task in cheminformatics(25). Most fingerprinting methods, however, are developed for pairwise molecule comparisons, e.g. for database retrieval or ligand screening, and are not so well suited for ML prediction and interpretability. Compared to molecular fingerprints, chemical ontologies, such as ChemOnt, actually offer interpretable hierarchy classes, and can be more discriminative of closely related molecules, such as natural products of the same family.

To predict the ChemOnt classes for a putative BGC product using only genomic features, we developed a novel ML method, called CHAMOIS (Chemical Hierarchy Approximation for secondary Metabolism clusters Obtained In Silico), which is trained on ChemOnt annotations of known BGC-derived compounds from MIBiG 3.1 (N=1,598 bacterial BGC corresponding to a total of N=1,034 ChemOnt classes, see Fig. 1 and *Sequence dataset preparation* section of the Methods). Instead of manual annotation, we used ClassyFire(24), a tool to automatically assign any molecule with a known chemical structure to the ChemOnt ontology. The hierarchical ChemOnt class annotation was subsequently treated as a binary vector (ones indicating class memberships) and for simplicity and scalability approached as a series of (independent) binary classification tasks. For reasons of interpretability, we opted for LASSO logistic regression as binary classification models(26). We excluded classes occurring in less than 5 compound groups, to enable evaluation with 5-fold stratified grouped cross-validation (*vide infra*). The remaining classes (N=539) were used to train LASSO classifiers (see *Compound dataset preparation* section of the Methods).

**Figure 1:**
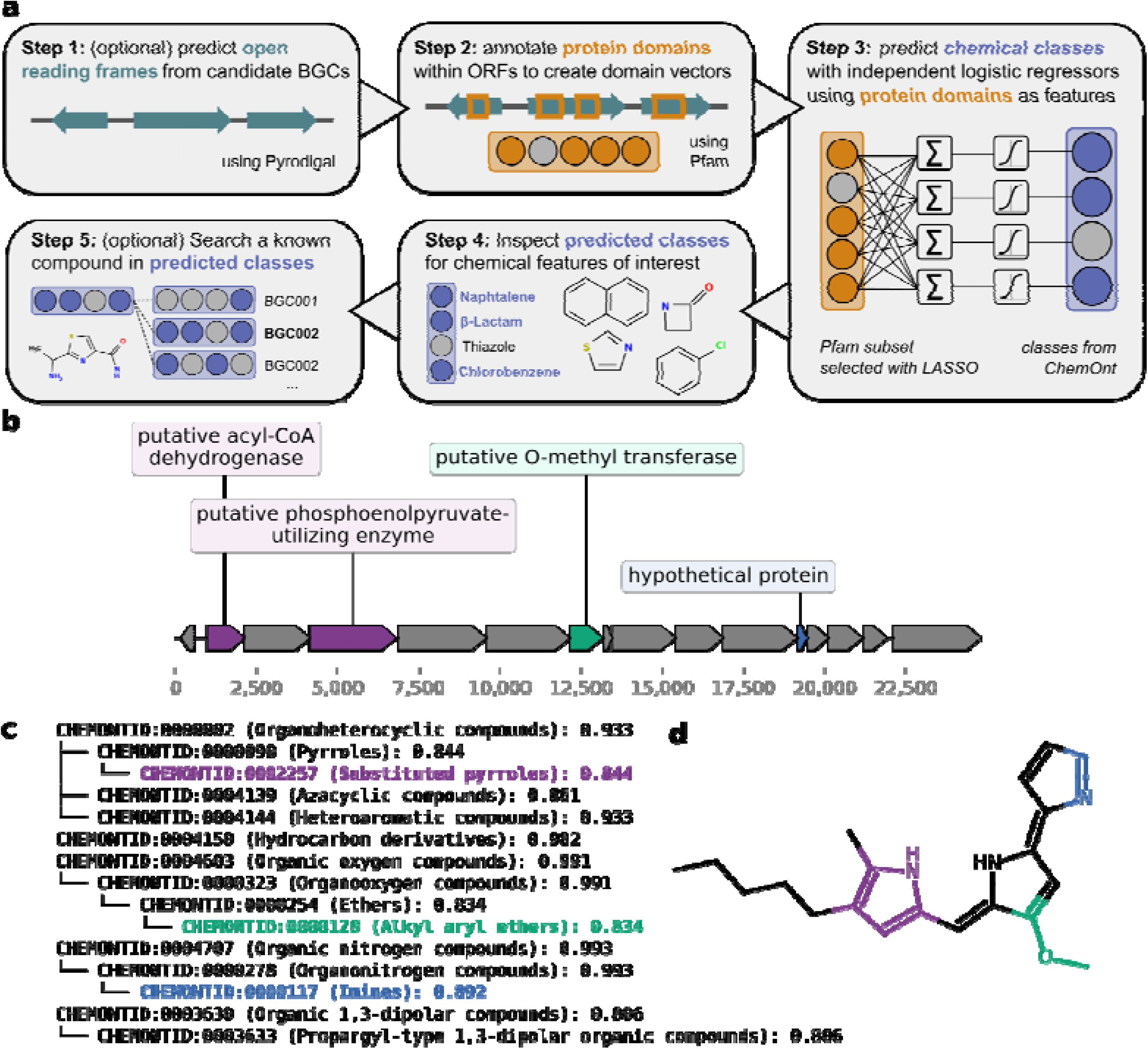
CHAMOIS, a novel method for predicting chemical properties of secondary metabolites from biosynthetic gene clusters (BGCs). **(a)** Workflow depiction of the chemical ontology prediction as implemented in CHAMOIS. First, CHAMOIS identifies the open reading frames in the input BGCs. Then, protein domains are annotated in the resulting ORFs using a subset of profile Hidden Markov Models from Pfam (N=896) selected as informative for CHAMOIS modeling. Each BGC is then encoded as a boolean feature vector indicating presence/absence of these Pfam domains. From these feature vectors, a multilabel logistic regression classifier predicts classes in the ChemOnt hierarchy (N=539). Resulting predictions can be inspected for classes of interest, or compared for similarity against a query compound. **(b)** Example of a genomic locus and **(c)** CHAMOIS prediction for the prodigiosin BGC of *Serratia marcescens* (BGC0000259)(96). The predicted classes are depicted as hierarchy alongside their posterior probabilities. CHAMOIS accurately predicted the Substituted pyrroles (CHEMONTID:0002257), Alkyl aryl ethers (CHEMONTID:0000128) and Imines (CHEMONTID:0000117) classes, highlighted on **(d)** the prodigiosin molecule in purple, green and blue respectively, with the PiKAChU(97) library. Genes containing domains that contributed to these class predictions are highlighted in the corresponding colour. CHAMOIS can produce an equivalent summary in tabular format with the `chamois explain cluster` command.

To prepare genomic sequence-derived features for training these classifiers, we first identified open reading frames (ORFs) for each BGC region(27,28), translated these and annotated them with Pfam 38.0 domains(29,30). Pfam is a curated database of protein domains which offers profile Hidden Markov Models (pHMMs) for automated domain annotation. This approach has been repeatedly shown to yield informative features for subsequent ML modeling(9,10,31). Here, we restricted Pfam feature space to domains that appeared in at least one BGC in MIBiG 3.1 resulting in binary feature vectors (of 3,334 dimensions) encoding presence/absence of each domain per BGC. On average, 26.5 unique Pfam domains were identified in the MIBiG 3.1 BGCs (Supplementary Fig. 1). Using Pfam domains facilitated interpretability, as LASSO models learnt to associate specific Pfam domains with compound classes (see section *CHAMOIS infers the biosynthetic capabilities of protein domains* of the Results). To train the classifiers, we excluded features appearing in less than 5 compound groups. The binary feature vectors with the remaining domains (N=927) were passed to every independent logistic regression classifier.

### Evaluating prediction of chemical properties

Since we decomposed the hierarchical topology of the ChemOnt class labels into binary classification tasks, we could separately evaluate each classifier using cross-validation. As MIBiG contains homologous clusters for the same compound across different species (e.g. coformycin(32), BGC0002039-40), as well as groups of closely related BGCs producing compounds of the same family (e.g kanamycin and tobramycin(33)), a naively sampled cross-validation would lead to biased accuracy estimates that are inflated by highly similar BGC-metabolite instances. To assess if CHAMOIS would be able to generalize beyond very similar instances, we first grouped BGCs based on the similarity of their produced compound, using Hamming distance(34) between MHFP6 fingerprints(25) to measure the distance of produced metabolites in chemical space (see *Compound dataset preparation* section of the Methods). We then used these groups (where all BGCs/compounds with pairwise chemical similarity of 0.5 and greater would be assigned to the same fold, total N=1,247 examples used for cross-validation) to perform stratified cross-validation preserving these groups, using 5 distinct folds for each of the 539 classifiers (see *Cross-validation* section of the Methods).

The median area under the receiver-operating characteristic (ROC) curve (AUROC) across all evaluated classes was 0.79 (Supplementary Table 1). However, because of the imbalance in the predicted classes (Supplementary Fig. 2a), AUROC values can be misleadingly high despite the underlying classifier having very low precision (i.e. having excess false-positives among the instances it predicts to belong to a given class). As an evaluation metric better reflecting classifier precision for these unbalanced tasks, we focused on the area under the precision-recall curve (AUPRC).

In our cross-validation, 120 ChemOnt classes were found to be predicted by CHAMOIS with an AUPRC of 0.5 or higher (Fig. 2a, see also Supplementary Table 1 for the metrics for each class). This constitutes an assessment of generalization across chemically dissimilar metabolites due to the grouping of similar metabolites in cross-validation folds (Fig. 2b). We also evaluated the performance of Random Forest classifiers in place of LASSO logistic regressions and found no significant differences in overall performance (Supplementary Fig. 3).

**Figure 2:**
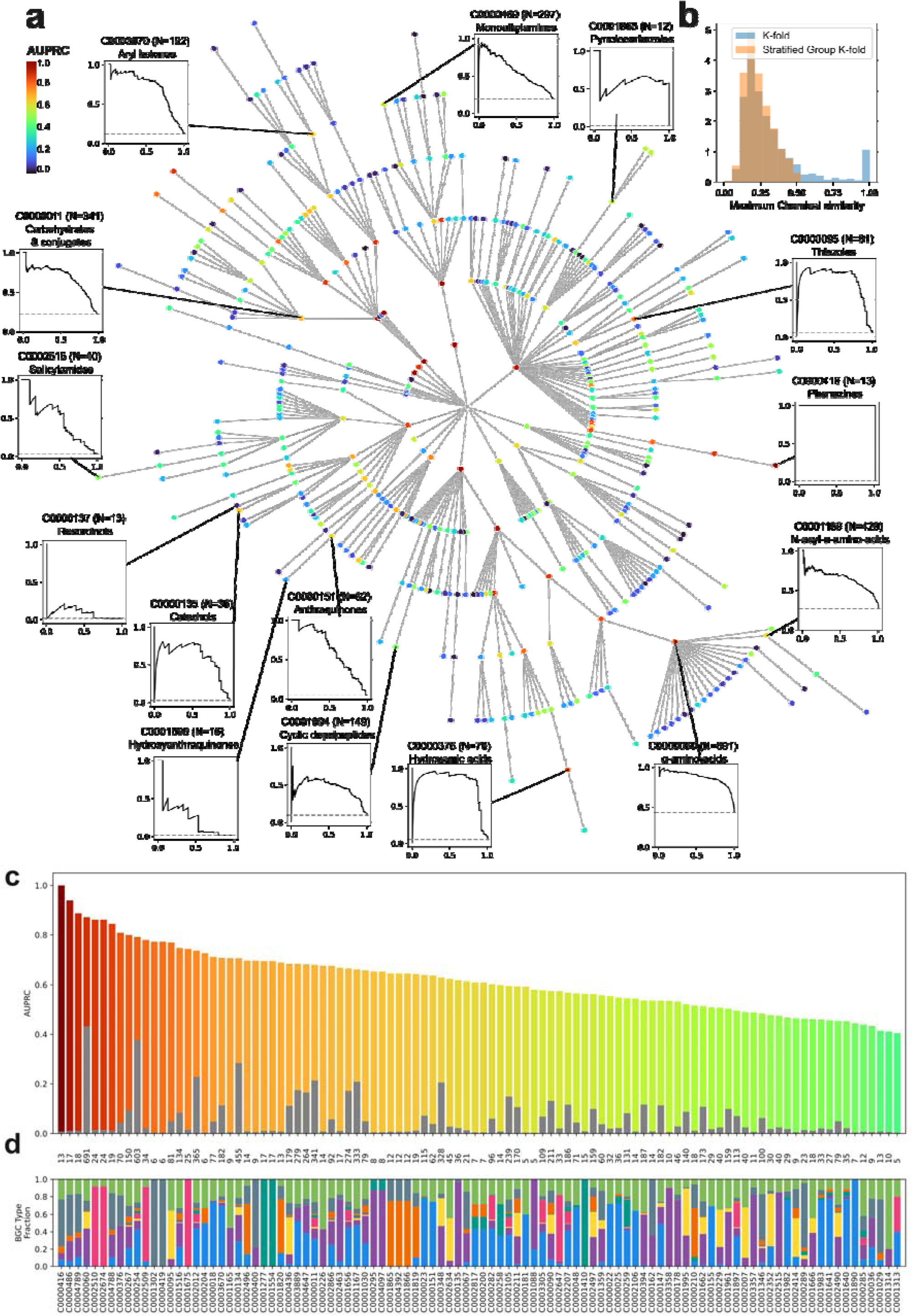
Evaluation of CHAMOIS against N=1,598 BGCs with known metabolites from MIBiG 3.1. **(a)** Model performance on every predicted ChemOnt class (N=539) using stratified grouped 5-fold cross-validation. Classes are displayed as nodes within the ChemOnt hierarchy and coloured according to CHAMOIS’s performance for the respective class, as assessed by the area under the precision-recall curve (AUPRC, see colour key). Precision-recall curves for selected classes are shown as insets against the baseline obtained by random guessing (dashed horizontal lines corresponding to class proportions). An interactive version of this panel can be found as part of CHAMOIS’ documentation at https://chamois.readthedocs.io/en/v0.2.0/figures/cv.html. **(b)** The barplot on the top-right corner indicates the distribution of chemical similarities (Hamming distance of MHFP6 fingerprints) between cross-validation folds, using either a naive 5-fold cross validation (blue) or a stratified cross-validation in which groups of similar compounds are kept in the same fold to assess generalization across distinct compounds (orange, used here with a distance cutoff of 0.5). **(c)** AUPRC values and for the 100 ChemOnt classes with the highest difference between CHAMOIS classifier AUPRC and random-guessing baseline AUPRC, shown as the coloured and grey section of each bar, respectively, sorted and coloured according to CHAMOIS AUPRC. Overall, predictors with >0.5 AUPRC were obtained for a total of 120 ChemOnt classes (Supplementary Table 1). **(d)** Distribution of BGCs in each ChemOnt class grouped by MIBiG types: *blue* Polyketide, *purple* NRP, *yellow* RiPP, *orange* Alkaloid, *pink* Saccharide, *green* Mixed (mostly Polyketide/NRP), *dark gray* Other.

Among the classes with the highest AUPRC were some high-level classes represented by many training instances, such as Organonitrogens (CHEMONTID:0000278, AUPRC=0.979, N_pos_=1,284) or Organoheterocyclic compounds (CHEMONTID:0000002, AUPRC=0.937 N_pos_=1,325). However, the class with the highest AUPRC was Phenazines (CHEMONTID:0000416, AUPRC=1.0, N_pos_=13), a rare class of tricyclic compounds known to be synthesised by a pathway of 7 conserved genes, *phzA-G*(35). In addition, CHAMOIS exhibited good performance on other relatively rare classes (Fig. 2c-d, Supplementary Fig. 2b): Cyclohexylamines (CHEMONTID:0002674, AUPRC=0.861, N_pos_=24), Thiazoles (CHEMONTID:0000095, AUPRC=0.769, N_pos_=81), Organohalogens (CHEMONTID:0000267, AUPRC=0.800, N_pos_=150), or Hydroxamic acids (CHEMONTID:0000376, AUPRC=0.832, N_pos_=71). Furthermore, the classes best predicted by CHAMOIS correspond to classes found across various types of BGCs, demonstrating that CHAMOIS predictions do not simply correlate with broad BGC types (Fig. 2d).

### CHAMOIS infers the biosynthetic capabilities of protein domains

CHAMOIS’s ML model is entirely data driven and unaware of any knowledge of natural product biosynthesis. Because we used LASSO classifiers, the learnt coefficients are sparse, which reduces the risk of overfitting and facilitates interpretation. To summarise the importance of few decisive domains in ChemOnt class prediction as learned by CHAMOIS, we extracted the 2 domains with the highest weight for every class, and used these relations to build a bipartite network capturing the strongest associations between ChemOnt classes and Pfam domains (Fig. 3). While each ChemOnt class could be predicted by a combination of many domains (on average 63, with a median of 34, domains received a non-zero LASSO coefficient), CHAMOIS classifiers for many ChemOnt classes indeed assigned a weight greater than 2.0 to a single domain (N=201, ∼37%) or two domains (N=106, ∼20%) (Supplementary Fig. 4, see Supplementary Table 2 for the complete matrix of weights learned by CHAMOIS).

**Figure 3:**
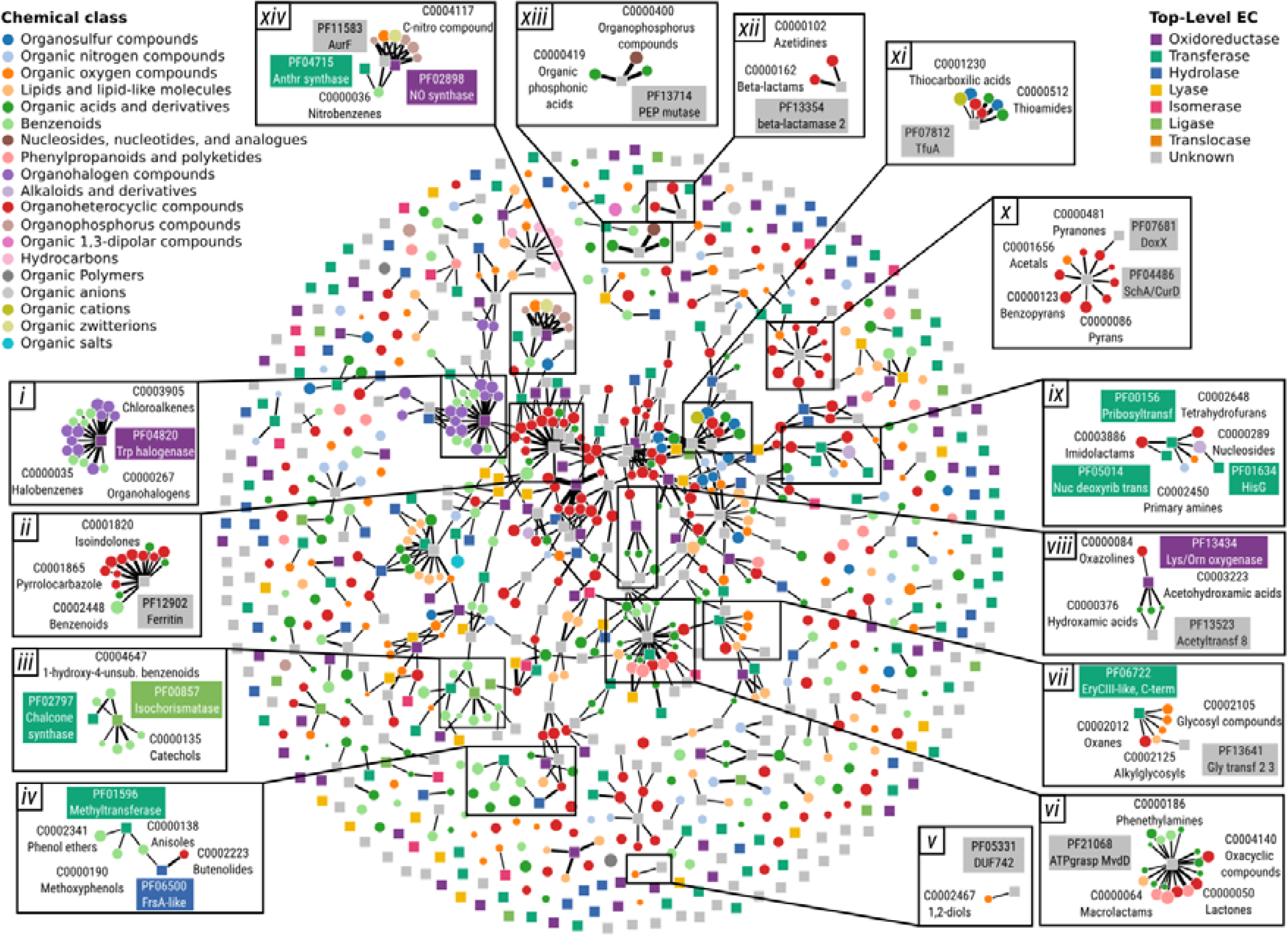
Network visualisation of associations between protein domains and chemical classes extracted from CHAMOIS’s logistic regression model coefficients (Supplementary Table 2). Each class (circle) is linked to Pfam domains (square) with weights >=2.0. Classes are coloured according to their highest-level ancestor in the ChemOnt taxonomy. Pfam domains are coloured by their top-level EC number, obtained with the ECDomainMiner tool(51). Several subgraphs feature associations supported by the literature (insets *i* to *iv, vi* to *ix*, and *xi* to *xiv)*, but also associations suggesting novel functions of uncharacterized domains (inset *v* and *x*, Table 1). An interactive version of this panel can be found as part of CHAMOIS’ documentation at https://chamois.readthedocs.io/en/v0.2.0/figures/network.html.

**Table 1:**
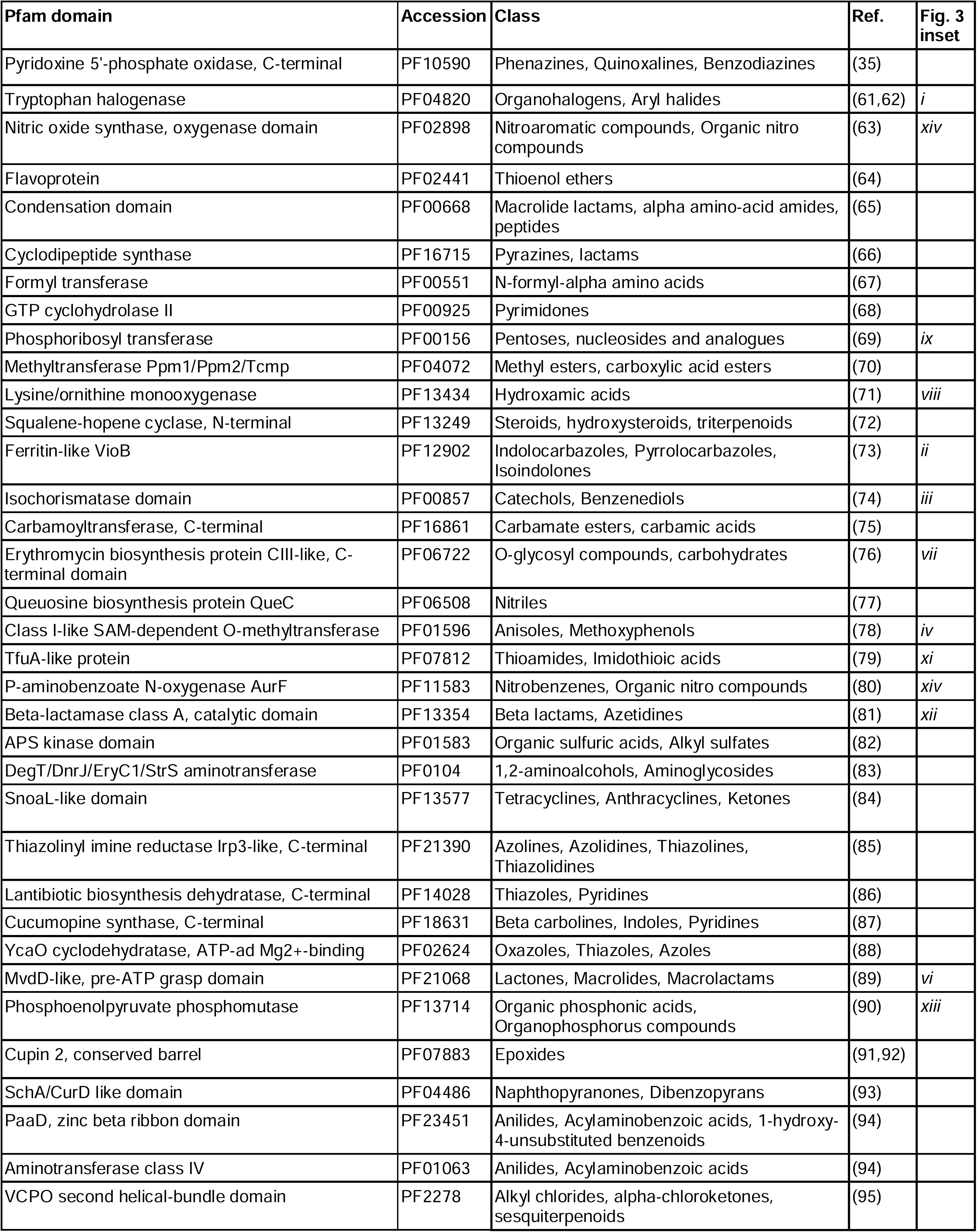
Select subset of highly weighted protein domain / chemical class associations learnt by CHAMOIS. Literature references consistent with CHAMOIS-inferred domain-class associations are included. Corresponding insets in Fig. 3 are indicated in the last column. An extended version of this table can be found in Supplementary Table 3.

In CHAMOIS’s classification model, a domain receiving a high weight suggests that it contributes to the formation of metabolite features that are characteristic of a given class. Without prior knowledge, CHAMOIS was able to recover known biosynthetic capabilities of certain Pfam domains. Among these examples of known biosynthetic domains were the Tryptophan halogenase domain (PF04820) linked to Aryl halides (CHEMONTID:0002866) biosynthesis, or the P-aminobenzoate N-oxygenase domain (PF11583) linked to Nitrobenzenes (CHEMONTID:0000036) biosynthesis (for 20 examples see Table 1 and Supplementary Table 3). This proof of principle that CHAMOIS extracted meaningful information about biosynthetic domains, prompted us to explore domain/metabolite class associations more broadly.

### CHAMOIS gives insight into the function of unknown domains

To identify the potential involvement of uncharacterized Pfam domains in the biosynthesis of certain compound classes, we collected a subset of Pfam domains devoid of functional annotation (as per InterPro 107.0(36)). In our dataset derived from MIBiG 3.1 an overwhelming majority of the BGC-associated domains still lack functional annotations: 1,428 of 1,576 BGCs (∼91%) contain at least one unannotated domain, also as a consequence of recent Pfam releases introducing new domains, such as DUF6069 (PF19545) present in 14 PKS BGCs, but still lack detailed annotations of their biosynthetic functions. From the set of domains with unknown function, we retained those to which CHAMOIS assigned a high weight (>2.0) in any ChemOnt classifier. This screening resulted in a list of 106 uncharacterized domains for which CHAMOIS inferred biosynthetic functions, for a total of 292 domain/class pairs (Supplementary Table 4, see *Identification of uncharacterized domains* section of the Methods) representing a rich resource for the discovery of biosynthetic functions.

While by construction, biosynthetic functional annotations could not be automatically retrieved for any of these domains, targeted literature searches recovered several of them to be well-described as such. For instance, the YcaO cyclodehydratase domain (PF02624) has been extensively studied for its involvement in post-translational modification of ribosomal peptides(37), yet it did not have any functional annotation in the corresponding InterPro entry (IPR003776). The associated ChemOnt classes in CHAMOIS were consistent with the literature, with a high weight assigned by the Thiazoles (CHEMONTID:0000095, w=4.005) classifier (Table 1, Supplementary Table 4). Other domains lacking annotations in the databases, but for which our approach recovered a function consistent with the scientific literature, included the Thiazolinyl imine reductase Irp3-like domain (PF21390), the Nitroreductase domain (PF00881), and the Lantibiotic biosynthesis dehydratase C-terminal domain (PF14028) (Table 1).

Beyond these confirmatory results, CHAMOIS also made novel predictions for which no direct literature support could be found, e.g. by connecting domains with truly unknown functions to a whole branch of the ChemOnt hierarchy. For instance, the Conserved hypothetical protein 95 (PF03602) received a high weight from the Dialkylarylamines (CHEMONTID:0003901, w=2.858, P=1.8E-3) and Tertiary alkylarylamines (CHEMONTID:0002454, w=2.784, P=1.3E-3) classifiers, hinting at a putative *N,N*-dimethyltransferase function. Among domains of unknown function *stricto sensu*, DUF742 (PF05331) received a high weight from the 1,2-diols (CHEMONTID:0002467, w=2.132, P=0.04) classifier, DUF4135 (PF13575) from Hydroxy acids and derivatives (CHEMONTID:0000472, w=2.392, P=6.9E-3) classifiers, DUF5837 (PF19155) from the Thiazolecarboxylic acids and derivatives (CHEMONTID:0002007, w=3.515, P=1.91E-10) classifier, and DUF6531 (PF20148) from the Hydroxypyridines (CHEMONTID:0004151, w=2.203, P=1.1E-3) classifier. Altogether these results suggest that CHAMOIS is capable of learning relevant associations between chemical classes and protein domains highlighting the promise of exploring these associations for the functional elucidation of uncharacterized biosynthetic domains.

### CHAMOIS outperforms PRISM 4.0 for structure prediction of Polyketides and NRPs

PRISM(23) is a state-of-the-art method for prediction of BGCs and their putative metabolites using a mixture of rules and *in silico* combinatorial generation of chemical structures. To compare CHAMOIS and PRISM 4 performance, we used the set of 1,279 BGCs known as the “gold standard BGCs”(38). We applied CHAMOIS to predict ChemOnt classes for these BGCs, and then searched the Natural Product Atlas(39) for the entry most similar to each CHAMOIS prediction. As done in Skinnider *et al.*, we computed the Morgan fingerprint of each prediction, and then took the median of the Tanimoto similarity to the true compound for each cluster (Supplementary Table 5, see the *Comparison to PRISM 4* section of the Methods). In this evaluation, CHAMOIS significantly outperformed PRISM 4.0 on structure prediction for BGCs encoding Polyketide or NRPs, while PRISM 4.0 slightly outperformed CHAMOIS on RiPP BGCs (Supplementary Fig. 5).

### CHAMOIS can prioritize putative BGCs for experimental characterization

A common task encountered by natural product biochemists is the identification of the producing cluster in the genome of a producer strain from which a metabolite of interest has already been identified. To pinpoint the biosynthetic machinery producing this metabolite, researchers typically first analyze the genome sequence with BGC prediction tools such as antiSMASH or GECCO, and subsequently manually inspect all predicted BGCs to identify the producing cluster. This search can be supported by CHAMOIS by predicting a ChemOnt fingerprint for each BGC and then quantifying its similarity to the fingerprint of the compound of interest, identifying the BGC with the highest likelihood of producing a certain metabolite.

To benchmark CHAMOIS’s ability to screen BGC predictions for a known compound, we assembled a test dataset of 70 experimentally-annotated bacterial BGCs which had been identified in a complete bacterial genome (Fig. 4a, Supplementary Table 6) and most of which showed moderate to no homology to any BGC from MIBiG 3.1, on which CHAMOIS was trained. The known metabolites were analysed with ClassyFire, as similarly done in the training dataset preparation to obtain a ChemOnt class vector. The source genomes were analyzed with antiSMASH and GECCO, to generate sets of candidate BGCs. In case of overlapping BGC predictions (made with different BGC finding tools), consensus clusters were obtained by taking the intersection gene set across overlapping predictions; when neither tool predicted a region overlapping the experimentally-validated BGC, it was manually added to the predictions. This resulted in 2,331 unique BGC predictions across all genomes (on average 37, ranging from 5 to 76 per genome). We then applied CHAMOIS to each BGC to predict a fingerprint consisting of the 530 ChemOnt classes covered by the model. For each genome, we calculated the similarity between all corresponding BGC fingerprints and that of the true compound using a probabilistic Jaccard index(40). Although most compounds in this benchmarking dataset belong to the Polyketide and NRP classes, the dataset represents an otherwise diverse set in the space of ChemOnt labels (Fig. 4b, see *Contextualization of benchmark data* section of the Methods).

**Figure 4:**
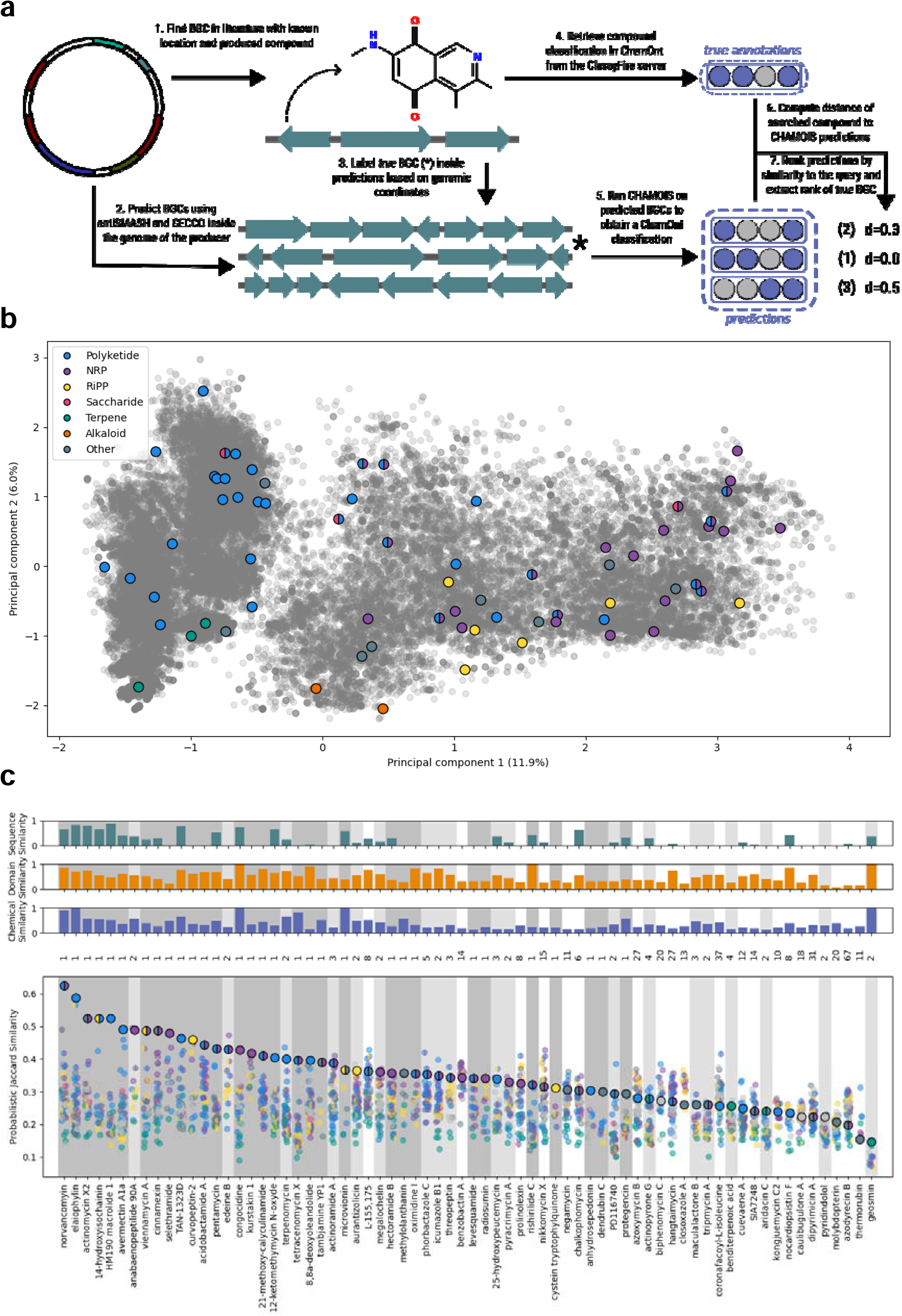
CHAMOIS can screen BGCs from within a given producer genome to pinpoint the cluster producing a given metabolite. **(a)** Graphical depiction of the prediction and evaluation framework. Shortly, as ground truth, experimentally validated BGCs with a known compound and found in a complete genome were extracted from literature (N=67). The known compounds were annotated with ClassyFire to obtain their reference ChemOnt classification. The source genomes were annotated for putative BGCs using antiSMASH (v7.0) and GECCO (v0.9.10) (see Methods). BGC predictions (N=2,331) were then passed to CHAMOIS to predict their ChemOnt classification. For each true compound, all predictions for a given genome were ranked using Probabilistic Jaccard similarity(40) between the predicted ChemOnt classes and the reference ChemOnt classification. **(b)** Principal component analysis (PCA) of ChemOnt class fingerprints highlighting compound diversity of the examples included in this benchmark (see *Contextualization of benchmark data* section of the Methods). NPAtlas compounds(39) are shown as gray background, while the compounds used for benchmarking are coloured by MIBiG classes. **(c)** Evaluation of CHAMOIS predictions using Probabilistic Jaccard similarity (y-axis) between predicted and true compounds for each producer genome (labeled on the x-axis). Each dot represents a BGC prediction, coloured by GECCO or antiSMASH predicted type (see colour code in (b)). The BGC producing the true compound is highlighted with an enlarged circle, and its rank among the predictions indicated above. A dark grey background indicates molecules where CHAMOIS correctly ranked the producing BGC with the highest similarity to the query (N=35, ∼52%), light-grey backgrounds highlight instances where the true compound ranked among the top five predictions (together N=50, ∼75%). The ranking produced by CHAMOIS is highly significant (P=1e-5, empirical permutation test with N=1e5 randomized rankings, see *BGC ranking significance* section of the Methods). The two barplots on top indicate sequence similarity (teal) and chemical similarity (lavender) to the closest examples in MIBiG 3.1 (used to train CHAMOIS) assessed by average nucleotide identity and Hamming distance between MHFP6 fingerprints, respectively (see *Establishing dissimilarity between benchmark and training sets* section of the Methods).

In this benchmark, CHAMOIS identified the correct cluster in 31 of the 70 cases (∼44%). In an additional 19 instances (∼27%), it ranked the true BGC among the top 5 clusters. That CHAMOIS in nearly three quarters of the benchmarking examples ranked the correct BGC very highly appears remarkable given that many of the genomes in this benchmark contained a large number of predicted BGCs of the same biosynthetic type and that fingerprint predictions by CHAMOIS often show only moderate similarity to that of the produced compound (<0.6, except for one BGC, Fig. 4c). Moreover, most of the examples used for benchmarking here share low chemical or sequence similarity with any secondary metabolites or BGC sequences contained in MIBiG 3.1 (used to train CHAMOIS), which underlines CHAMOIS’s capacity to generalize to diverse cluster architectures and metabolite classes. For instance, CHAMOIS successfully recovered the BGC for 8,8a-deoxyoleandolide across 64 predicted loci despite the cognate BGC only sharing ∼2% identity with the closest MIBiG BGC (BGC0000898, desosamine). Overall these results suggest that CHAMOIS could be a useful tool to expedite the identification of the BGC corresponding to interesting metabolites in isolated producer strains or low-complexity metagenomes.

## Discussion

Predicting secondary metabolites from their cognate BGCs *in silico* is an important problem, the urgency and potential of which is highlighted by the growing divide between available sequence and metabolite data. While previous research has resulted in prediction tools for certain BGC types, CHAMOIS is the first-of-its-kind universal open-source machine-learning method for predicting the ontological classification of a BGC product from its genomic sequence. In our study we carefully assessed its predictive capacity across dissimilar compound subsets in cross-validation to avoid overoptimistic performance reports (Fig. 2). In addition we performed a benchmark on an additional external dataset, which mostly contained BGCs and metabolites without similarity to the training data (Fig. 4). A limitation of our method, which implicitly relies on protein domains involved in biosynthesis, is the limited prediction performance for RiPP BGC products due to several RiPP-specific challenges. They often produce large molecules, the complexity of which can sometimes not be fully captured by ClassyFire. As an additional challenge, the structure of RiPP molecules largely depends on the precursor peptide sequence, not only on tailoring enzymes; while the latter can be captured by Pfam annotations, the precursor peptide typically remains unannotated. Even so, as CHAMOIS allows the prediction of tailoring enzyme modifications, it may potentially be useful in conjunction with type-specific methods that predict a BGC compound backbone, such as RiPPMiner(21).

CHAMOIS is an entirely data-driven method using multilabel LASSO logistic regression to infer ChemOnt classes. While LASSO regularisation avoids overfitting and enforces sparse weights useful for feature selection, it however comes with a drawback in the case of co-occurring features, where it will often arbitrarily select one and disregard the others(41). From a biological point of view, co-occurring features may often be found in BGCs encoding enzyme complexes, which require the presence of more than one protein to catalyse a given reaction. In these cases, LASSO-derived associations between domains and ChemOnt classes will on the one hand tend to capture only one complex member, so that the others remain undetected. On the other hand false-positive predictions may occur when an incomplete pathway is encountered. On the upside, CHAMOIS’s LASSO classifiers are fast to retrain and evaluate, and will easily cope with – and benefit from – more data becoming available in future BGC database updates. Likewise, the inclusion of new domain features will be straightforward as CHAMOIS does not rely on their functional annotation. In particular, when more natural products belonging to rare ChemOnt classes become available in the future, CHAMOIS will be capable of predicting a more diverse set of classes and will provide an automated approach to obtain clues as to the protein domains characteristic of these classes and their key biosynthetic reactions (Fig. 3). Eventually, an even larger amount of training data could support modeling more complex relationships between genes, e.g. using modern AI architectures such as Autoencoders in the future. However, as currently available data sets are limited in size, such methods are unlikely to outperform CHAMOIS’s more classical ML approach.

As CHAMOIS enables (inferential) translation of BGCs into a chemical space, it holds potential for exploring natural product space beyond what is feasible based on genomic representations alone. This translation can for instance be used to scan sets of BGC predictions for members of a chemical class of interest, or to cluster BGCs based on their putative product properties, or to assess biosynthetic diversity in terms of not only (meta-)genomic sequence similarity, but also biochemical class representation. Additionally, the genetic basis of this chemical space (in terms of biosynthetic protein domains predicted by CHAMOIS) can be explored as an additional layer of information to guide the discovery of new enzymatic domains in well-characterized as well as newly discovered BGCs.

## Conclusion

Chamois represents a scalable, entirely data-driven computational method for inferring secondary metabolite properties from their cognate BGCs that is – unlike most previous methods – not restricted to certain types of clusters. It facilitates exploring the biochemical space of natural products on the basis of rapidly growing genomic BGC resources as an important step towards genotype-phenotype prediction for this highly relevant area of microbial metabolism.

## Methods

### Sequence dataset preparation

BGC sequences for MIBiG 3.1(42) entries were downloaded from the MIBiG website (https://mibig.secondarymetabolites.org) as GenBank files. 567 BGCs were filtered based on a manually curated list, to exclude eukaryotic records, and records with known annotation issues (Supplementary Table 7). The cluster sequences were trimmed to the coordinates of the outermost genes of the clusters. The coordinates of 82 clusters were programmatically corrected to retain only the biosynthetic core based on the reference literature (see data/scripts/mibig/download_records.py in the project repository). The BGC records were then annotated with the CHAMOIS CLI (chamois annotate). Cluster genes were called using Pyrodigal(28) v3.6.3 wrapping Prodigal(27) v2.6.3 in *meta* mode with closed ends (-c -p meta). The genes were then mapped against Pfam(43) v38.0 using hmmsearch as implemented in PyHMMER(30) v0.11.1 wrapping HMMER (http://hmmer.org) v3.4. The Pfam domains were filtered using the “trusted” bitscore cutoffs of each profile HMM (--cut_tc). From each BGC, a boolean feature vector of annotated Pfam domains (d=3,146) was constructed, which was stored in an HDF5 file using the anndata library(44) v0.12.3.

### Compound dataset preparation

The BGC metadata for MIBiG 3.1(42) and 4.0(7) entries were downloaded from the MIBiG website (https://mibig.secondarymetabolites.org) as JSON files, containing the structures of the compounds as SMILES. 110 cluster compounds were programmatically corrected for incorrect name or SMILES annotations (see the download script data/scripts/mibig/download_compounds.py in the project repository). For compounds missing a structure in the metadata, the corresponding NPAtlas structure was fetched if a cross-reference existed. When compounds contained no cross-reference, they were mapped by name against the NPAtlas(45), or, failing that, against PubChem(46) (https://pubchem.ncbi.nlm.nih.gov). All obtained SMILES strings were then converted into InChi using RDKit(47) v2023.9.6 (http://www.rdkit.org). The InChi strings of each compound were submitted to annotation on the ClassyFire(24) web server (http://classyfire.wishartlab.com). The classification of each compound was then encoded into a binary vector of ChemOnt classes for each compound and assigned to its cognate BGCs; for BGCs producing more than one compound, the compound with the highest number of ChemOnt classes was used. For MIBiG 3.1, 1,598 BGCs could be successfully labelled this way. The resulting labels were stored in an HDF5 file using the anndata library(44) v0.12.3. For further stratification, we grouped BGCs using a pairwise Hamming distance(34) cutoff of 0.5 between their MHFP6 fingerprints(25) computed using RDKit(47) v2023.9.6 (http://www.rdkit.org) resulting in 1,180 groups of BGCs that were distinct to each other with respect to the encoded metabolites.

### Model training

The training dataset was first filtered to remove clusters shorter than 1,000 bp. Classes with less than 5 positive or negative occurrences across groups were excluded from the label matrix. A LASSO logistic regression model was trained on each target label independently (N=539) with the LogisticRegression classifier implemented in scikit-learn(48) v1.7.2, using the LIBLINEAR(49) solver with L1 regularisation and 100 maximum iterations. Features that received zero weights (coefficients) from all classifiers were discarded, and only the remaining features were retained in the model (d=896). The weights were then combined into a single weight matrix and stored alongside the model metadata in JSON format. The whole training procedure is implemented in the train command of the chamois command-line tool.

### Cross-validation

The model was validated using a cross-validation strategy. First, classes with less than 5 positive or negative occurrences across groups were excluded from the label matrix. Then, for each class of the label matrix, a 5-fold cross-validation was run, using a StratifiedGroupKFold from scikit-learn(48) v1.7.2 to split the data according to the molecular-similarity groups built from MHFP6 fingerprints(25), found in the “group” column of the observation table of the dataset. The probabilities generated for each label were combined into a probability matrix, for which the area under the receiver-operator characteristic curve (AUROC) and area under the precision-recall curve (AUPRC) were computed with micro- and macro-averaging using the roc_auc_score and average_precision_score functions of scikit-learn respectively. The whole evaluation procedure is implemented in the cvi command of the chamois command-line tool.

### Visualization of cross-validation performance across the ChemOnt hierarchy

The pronto Python library (https://github.com/althonos/pronto) v2.6.0 was used to load the ChemOnt ontology from OBO format, and converting the class hierarchy into an adjacency matrix, treating each class as a node and each subclassing relationship as an edge in the tree. This tree was then displayed with the Vega visualization framework (https://vega.github.io/) in a radial tree layout, using AUPRC values computed with the average_precision_score as described above to color the nodes. Individual precision-recall curves were generated with matplotlib(50) v3.10.6 (Fig. 2).

### Network visualization

The weights of the trained model were extracted from the CHAMOIS LASSO classifiers (Supplementary Table 2). For each classifier corresponding to a ChemOnt class, the two domains with the highest weights were extracted to form a bipartite graph with chemical classes and protein domains as nodes. To categorize the Pfam domains, the predictions from the ECDomainMiner tool(51) were used to label domains by top-level Enzyme Commision (EC) number. To categorize ChemOnt classes, their top-level ancestor in the ChemOnt hierarchy was extracted. The graph was then displayed with the Vega visualization framework (https://vega.github.io/) using force-directed layout (Fig. 3).

### Identification of uncharacterized domains

Domain annotations were downloaded from InterPro(36) 107.0 in XML format (https://www.ebi.ac.uk/interpro/download/). We collected all Pfam(43) 38.0 domains, and retained those as “uncharacterized” which did not have a corresponding InterPro entry with an Enzyme Commission (EC) number or a Gene Ontology(52) term. We further analyzed domains that were assigned a weight of at least 2.0 by any classifier within CHAMOIS (Supplementary Tables 3 and 4).

### Significance of domain to chemical class associations

For each uncharacterized domain/chemical class pair extracted , a contingency table was built, counting occurrences inside the MIBiG 3.1 dataset. For each table, Fisher’s exact p-value was calculated using the scipy.stats.fisher_exact function of SciPy(53) v1.16.3. The p-values are reported both uncorrected and corrected with Bonferroni correction (Supplementary Table 4).

### Comparison to PRISM 4

The BGCs records of the 1,281 “Gold Standard” BGCs from PRISM were downloaded from Zenodo(38). The PRISM 4-predicted SMILES for each BGC were obtained from the Supplementary Material of the PRISM 4 paper(23). The ChemOnt classes for each BGCs were predicted with the chamois predict command using default parameters. The predicted ChemOnt classes were used to search the Natural Product Atlas(39) with the chamois search command using default parameters. For each cluster, the highest ranking compounds were selected (allowing ties) and their SMILES extracted as the predicted CHAMOIS structures. Using the evaluation code of Skinnider *et al.* (https://github.com/Adapsyn/prism-4-paper), we computed the Tanimoto coefficient of the ECFP6 fingerprints of the predicted molecules to the true compound for CHAMOIS as well as the methods present in the PRISM4 “Gold Standard” dataset: PRISM 1(54), PRISM 4, NP.searcher(55) and antiSMASH 4.0(56) (Supplementary Table 5). For BGCs where all methods successfully predicted a structure (n=373), the median Tanimoto coefficient was extracted, and summary boxplots were generated similarly to Fig. 2c and 2e of Skinnider *et al.* (Supplementary Fig. 5a-b). For all BGCs, we extracted the median Tanimoto of CHAMOIS and PRISM 4 predictions, and grouped them by MIBiG BGC types across PK, NRP, RiPP and Other BGCs, respectively, to allow direct comparison (Supplementary Fig. 5c). Significance was assessed using unpaired *t*-test computed with the scipy.stats.ttest_ind function of SciPy(53) v1.16.3.

### BGC screening in producer genomes

To evaluate how well CHAMOIS could pinpoint the cognate BGC for a given compound, we assembled a dataset of 70 experimentally-validated BGCs not included in MIBiG 3.1(42) that were contained in 65 (near-)complete bacterial genomes (Supplementary Table 6). Each of these genomes was newly annotated with the antiSMASH(6) web server v8.0 and GECCO(10). For regions where both tools predicted a BGC, their intersection was kept. When neither tool predicted a region overlapping the experimentally-validated BGC, it was manually added to the predictions. All predicted BGCs (N=2,527) were then annotated with CHAMOIS to predict a ChemOnt classification. For each validated BGC/compound pair, the distance between the true compound and all predictions was computed using Probabilistic Jaccard similarity(40) between their ChemOnt classes. The BGC predictions were then sorted and assigned a relative rank based on their similarity to the ChemOnt classes of the true compound.

### Contextualization of benchmark data

The NPAtlas v2024_03 entries(57) were downloaded from the NPAtlas(45) server. Each compound was binarized into an indicator label vector as described earlier (see *Compound dataset preparation*), using the pre-computed ChemOnt annotations provided by the NPAtlas. The complete indicator matrix was used to train a Principal Component Analysis (PCA) using the PCA class from scikit-learn(48) v1.7.2 with default parameters. The compounds from the benchmark dataset were labeled with the ClassyFire(24) web server (http://classyfire.wishartlab.com) as described earlier (see *Compound dataset preparation*). The compounds were then projected into the aforementioned principal component space. The resulting principal components were plotted with matplotlib(50) v3.10.0 (Fig. 4b).

### BGC ranking significance

To evaluate the significance of the ranking produced by CHAMOIS on the whole dataset, the relative rank of the true compound in the CHAMOIS predictions sorted by Probabilistic Jaccard similarity was computed using the rankdata function of SciPy(53) v1.15.2 (with parameters method=”dense”). An empirical background distribution was computed using randomization (N=100,000) by selecting a “true” BGC at random in each genome, and averaging their relative rank to the true compound. A lower relative rank than the one obtained by CHAMOIS was not found in any of the random permutations.

### Establishing dissimilarity between benchmark and training sets

We queried the BGC predictions intersecting the true cognate BGCs against MIBiG 3.1 BGCs using skani(58) v0.2.2 (with parameters -c10 -m50) wrapped in Pyskani(59) v0.1.3. For each hit, the query ANI was computed, where □□□□□ □□□ _ □□□□□□□□ _ □□□□□ □□□□□□□□ with □□□□□□□□ and □□□□□ □□□□□□□□ computed by skani. In addition, we measured the chemical similarity between the true compound and every MIBiG 3.1 BGC using Hamming distance between MHFP6 fingerprints. The highest query ANI and the highest chemical similarity for each BGC were plotted with matplotlib(50) v3.10.0 (Fig. 4c).

## Declaratations

### Ethics approval and consent to participate

Not applicable.

### Consent for publication

Not applicable.

## Availability of data and materials

### Code availability

CHAMOIS code is open source under the GNU General Public License 3.0 or later (GPL-3.0-or-later), publicly hosted in a git repository on GitHub at https://github.com/zellerlab/CHAMOIS. An archive of the version used for this study (v0.2.0) is available in the Zenodo repository (https://zenodo.org/records/17849623). CHAMOIS can be installed for Python 3.7 and later on UNIX operating systems (Linux, MacOS, Windows WSL, etc.) from the Python Package Index (PyPI; https://pypi.org/project/chamois-tool) and from the Bioconda(60) channel of the conda package manager (https://anaconda.org/bioconda/chamois). A self-contained Docker image with all dependencies can be obtained from the GitHub Container Registry (https://ghcr.io/zellerlab/chamois).

### Data availability

The datasets generated during the current study are available in the Zenodo repository (https://zenodo.org/records/17849853). CHAMOIS weights are included in this published article in Supplementary Table 2.

## Competing interests

The authors declare no competing interests.

## Funding

This work was supported by the European Molecular Biology Laboratory (EMBL); the SFB 1371 of the German Research Foundation (Deutsche Forschungsgemeinschaft, DFG) [395357507 to G.Z.], and a LUMC Fellowship [to G.Z.].

## Author’s contributions

Software development and computational analyses were performed by ML, with suggestions from GZ. ML and GZ conceived the study, designed the figures and co-wrote the manuscript.

## Acknowledgements

We are grateful to Joachim Hug and Michael Zimmerman (both EMBL Heidelberg), to Laura Carroll (Umeå University), to Justin van der Hooft and Marnix Medema (both Wageningen University) for their suggestions during development. We are indebted to the European Molecular Biology Laboratory (EMBL) and its IT Services Team for providing and administrating HPC resources.

## Notes

https://github.com/zellerlab/CHAMOIS

https://zenodo.org/records/17849853

https://zenodo.org/records/17849623

### Competing Interest Statement

The authors have declared no competing interest.

### Summary of Updates

Updated abstract; curated BGC datasets and re-generated features with Pfam 38.0; model re-trained on updated MIBiG 3.1 dataset; Updated Fig. 2a, Fig. 3 and Fig. 4c; Added Fig. 2d; Added supplemental figures; Added PRISM 4 datasets and performance comparison as supplemental figure; Updated Table 1; Improved Methods section; Revised overall manuscript text.

