## Supplementary Figures for "Machine learning inference of natural product chemistry across biosynthetic gene cluster types"

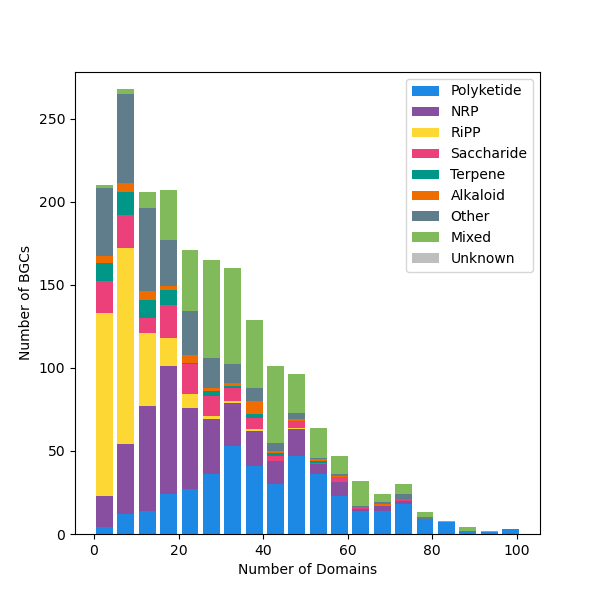

**Supplementary Figure 1**: Distribution of Pfam 38.0 in MIBiG 3.1 BGCs used for training CHAMOIS. All Pfam domains are included, prior to any feature selection. BGC type is shown according to MIBiG 3.1 metadata.

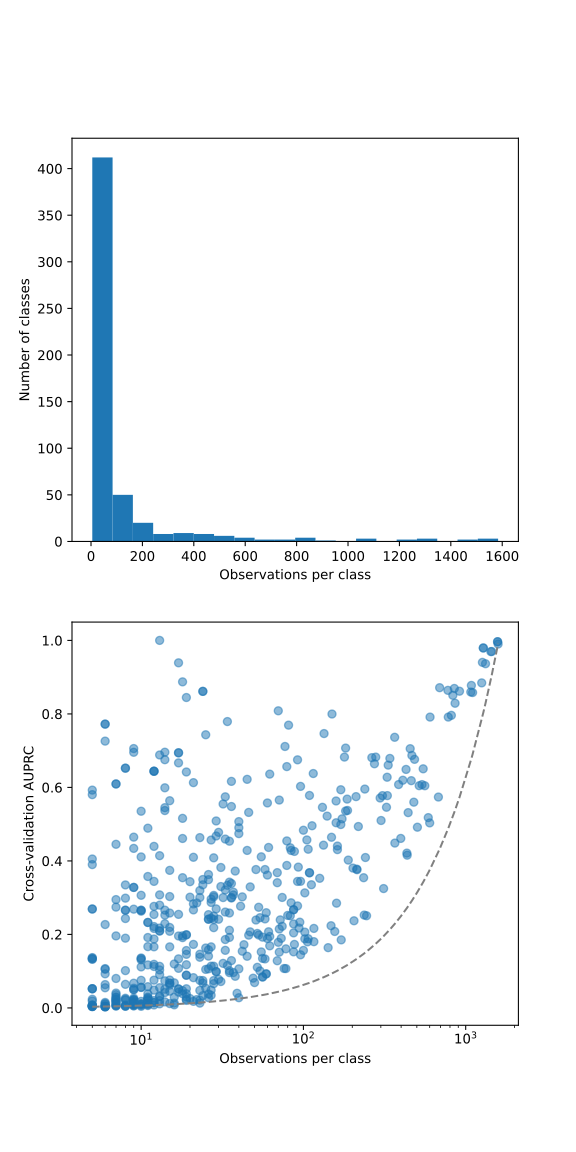

**a**

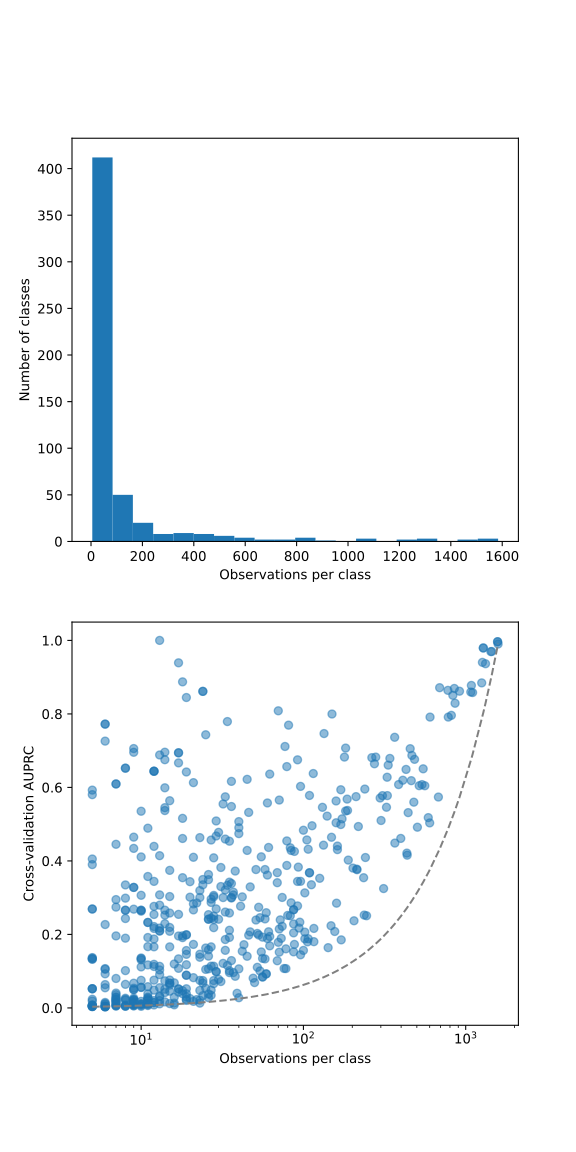

**b**

**Supplementary Figure 2**: ChemOnt classes distribution in MIBiG 3.1 BGCs. **(a)** Histogram of number of ChemOnt classes by number of observations (BGCs in MIBiG 3.1). **(b)** Performance of CHAMOIS measured by AUPRC in function of the number of positive observations per ChemOnt class.

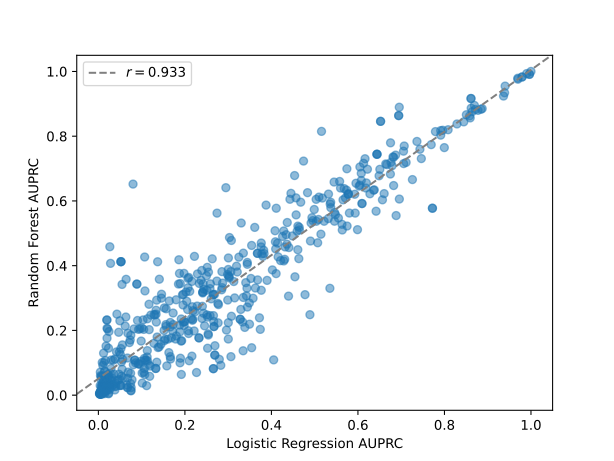

**Supplementary Figure 3**: Correlation between Logistic Regression and Random Forest classifier performances. Both models were trained and evaluated independently on the MIBiG 3.1 dataset and the AUPRC reported as a measure of performance.

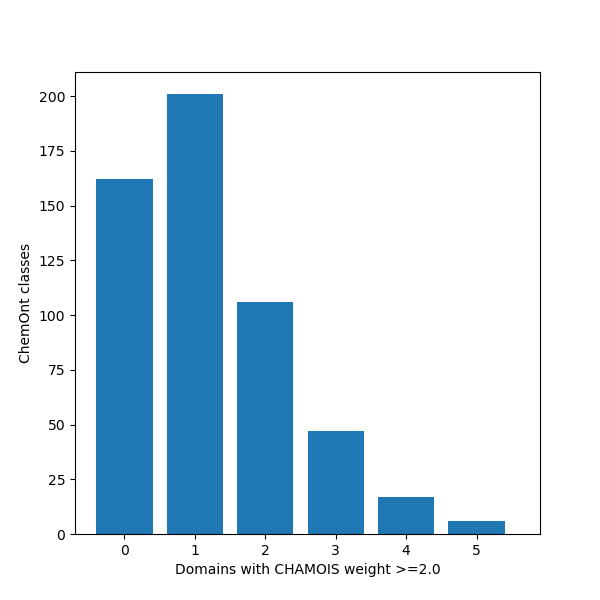

**Supplementary Figure 4**: Distribution of CHAMOIS models with high weight. For each CHAMOIS classifier, the number of features receiving a weight greater than 2.0 were extracted, similarly to the network drawn in Fig. 3.

**a**
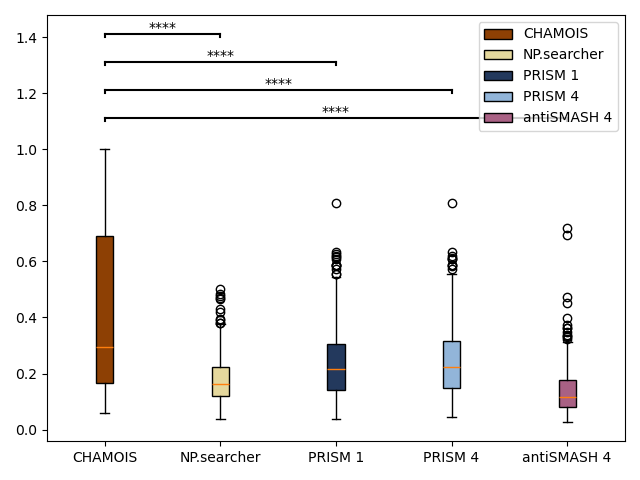

**b**
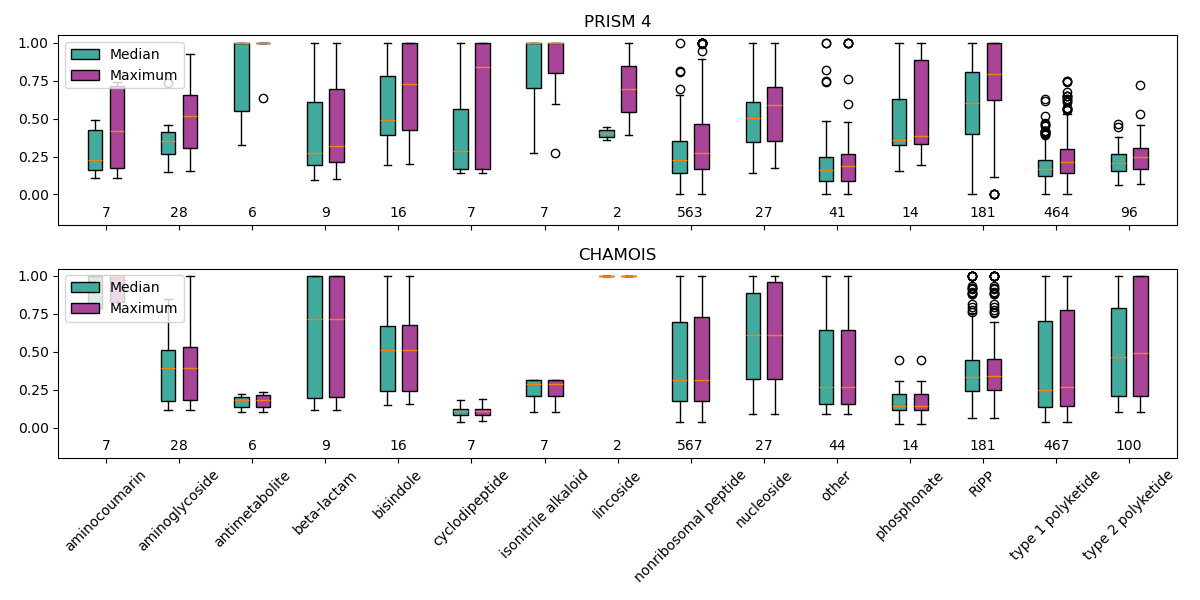

**c**

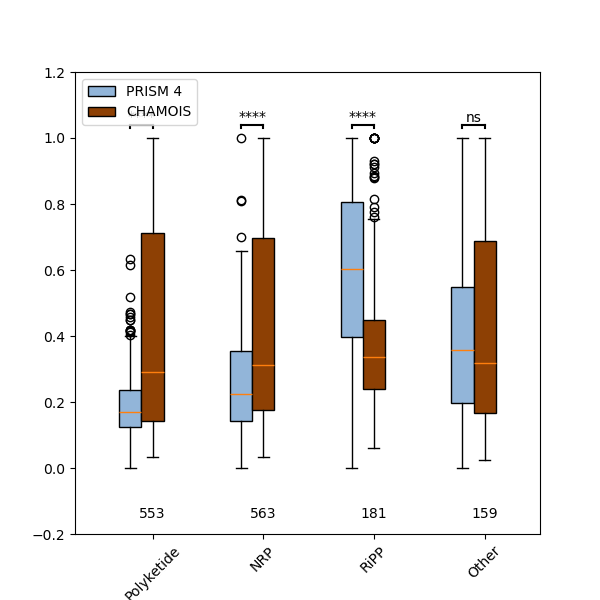

**Supplementary Figure 5**: Comparison of chemical prediction for unknown BGCs. **(a)** Median Tanimoto coefficient between true and predicted structure for the subset of BGCs from the PRISM “gold standard” dataset (https://zenodo.org/record/3985982) where all methods generated an output (N=385). **(b)** Median and maximum Tanimoto coefficients between true and predicted structures generated by CHAMOIS v0.2.0 (with NPAtlas 2024_09) and PRISM 4 for the “gold standard” set (N=1,281), grouped by biosynthetic family. **(c)** Median Tanimoto coefficieints between true and predicted structures generated by CHAMOIS v0.2.0 (with NPAtlas 2024_09) and PRISM 4 for the “gold standard” set (N=1,281), grouped by MIBiG type. Top, statistic significance of the comparison between methods (**** p<0.0001, *** p<0.001, ** p<0.01, * p<0.05, *ns* otherwise) given from an independent *t*-test. See *Comparison to PRISM 4* section of the Methods.
